# The QseB response regulator imparts tolerance to positively charged antibiotics by controlling metabolism and minor changes to LPS

**DOI:** 10.1101/2023.01.10.523522

**Authors:** Melanie N. Hurst, Connor J. Beebout, Alexis Hollingsworth, Kirsten R. Guckes, Alexandria Purcell, Tomas A. Bermudez, Diamond Williams, Seth A. Reasoner, M. Stephen Trent, Maria Hadjifrangiskou

## Abstract

The modification of lipopolysaccharide (LPS) in *Escherichia coli* and *Salmonella spp*. is primarily controlled by the two-component system PmrAB. LPS modification allows bacteria to avoid killing by positively charged antibiotics like polymyxin B. We previously demonstrated that in uropathogenic *E. coli* (UPEC), the sensor histidine kinase PmrB also activates a non-cognate transcription factor, QseB, and this activation somehow augments polymyxin B tolerance in UPEC. Here, we demonstrate – for the first time – that in the absence of the canonical LPS transcriptional regulator, PmrA, QseB can direct some modifications on the LPS. In agreement with this observation, transcriptional profiling analyses demonstrate regulatory overlaps between PmrA and QseB in terms of regulating LPS modification genes. However, both PmrA and QseB must be present for UPEC to mount robust tolerance to polymyxin B. Transcriptional and metabolomic analyses also reveal that QseB transcriptionally regulates the metabolism of glutamate and 2-oxoglutarate, which are consumed and produced during the modification of lipid A. We show that deletion of *qseB* alters glutamate levels in the bacterial cells. The *qseB* deletion mutant, which is susceptible to positively charged antibiotics, is rescued by exogenous addition of 2-oxoglutarate. These findings uncover a previously unknown mechanism of metabolic control of antibiotic tolerance that may be contributing to antibiotic treatment failure in the clinic.

**IMPORTANCE:** Although antibiotic prescriptions are guided by well-established susceptibility testing methods, antibiotic treatments oftentimes fail. The presented work is significant, because it uncovers a mechanism by which bacteria transiently avoid killing by antibiotics. This mechanism involves two closely related transcription factors, PmrA and QseB, which are conserved across Enterobacteriaceae. We demonstrate that PmrA and QseB share regulatory targets in lipid A modification pathway and prove that QseB can orchestrate modifications of lipid A in *E. coli* in the absence of PmrA. Finally, we show that QseB controls glutamate metabolism during the antibiotic response. These results suggest that rewiring of QseB-mediated metabolic genes can lead to stable antibiotic resistance in subpopulations within the host, thereby contributing to antibiotic treatment failure.

## INTRODUCTION

Antibiotic resistance is a global pandemic and includes high rates of antibiotic treatment failure. One in every ten antibiotic prescription fails even when the clinical laboratory’s antimicrobial susceptibility panel predicts susceptibility to a given drug. (1-5). The molecular underpinnings behind such treatment failures remain largely undefined. This work elucidates a previously uncharacterized mechanism in uropathogenic *Escherichia coli* (UPEC) that leads to transient tolerance to polymyxin B and other positively charged antibiotics.

Enterobacteriaceae are common human pathogens, accounting for urinary tract infections, bloodstream infections and pneumonias (6). Among the antibiotics currently used to treat infections caused by multi-drug resistant Enterobacteriaceae, are aminoglycosides and polymyxins, which are polycationic in nature and therefore contact the bacterial cell envelope by binding to negatively charged moieties on the lipopolysaccharide (LPS) (7, 8). This interaction leads to increased permeability and penetration of the aminoglycoside or polymyxin into the periplasm. A mechanism used by bacteria to repel cationic antibiotics is to make the bacterial cell envelope less negatively charged (8). Altering the net charge of the envelope can be accomplished through different mechanisms, including dephosphorylation of the lipid A component of LPS (**Figure 1A**), or addition of positively charged groups – such as phosphoethanolamine and aminoarabinose – directly to the lipid A group during synthesis (9). In *E. coli*, the majority of lipid A modifications occur during LPS biogenesis (**Figure 1A**) at the periplasmic leaflet of the inner membrane (10). The ArnB transaminase catalyzes a reversible reaction of undecaprenyl-4-keto-pyranose to undecaprenyl 4-amino-4-deoxy-L-arabinose by consuming glutamate and producing oxoglutarate in the process. Early work by the Raetz group indicated that the ArnB-mediated addition of amino-arabinose is energetically unfavorable and requires excess glutamate, as determined through an *in vitro* radiometric enzymatic assay (11). Given the central role of glutamate in *E. coli* physiological functions (12-14), this raises a fundamental question of how *E. coli* manages the metabolic burden associated with modifications of the LPS.

**Figure 1.**
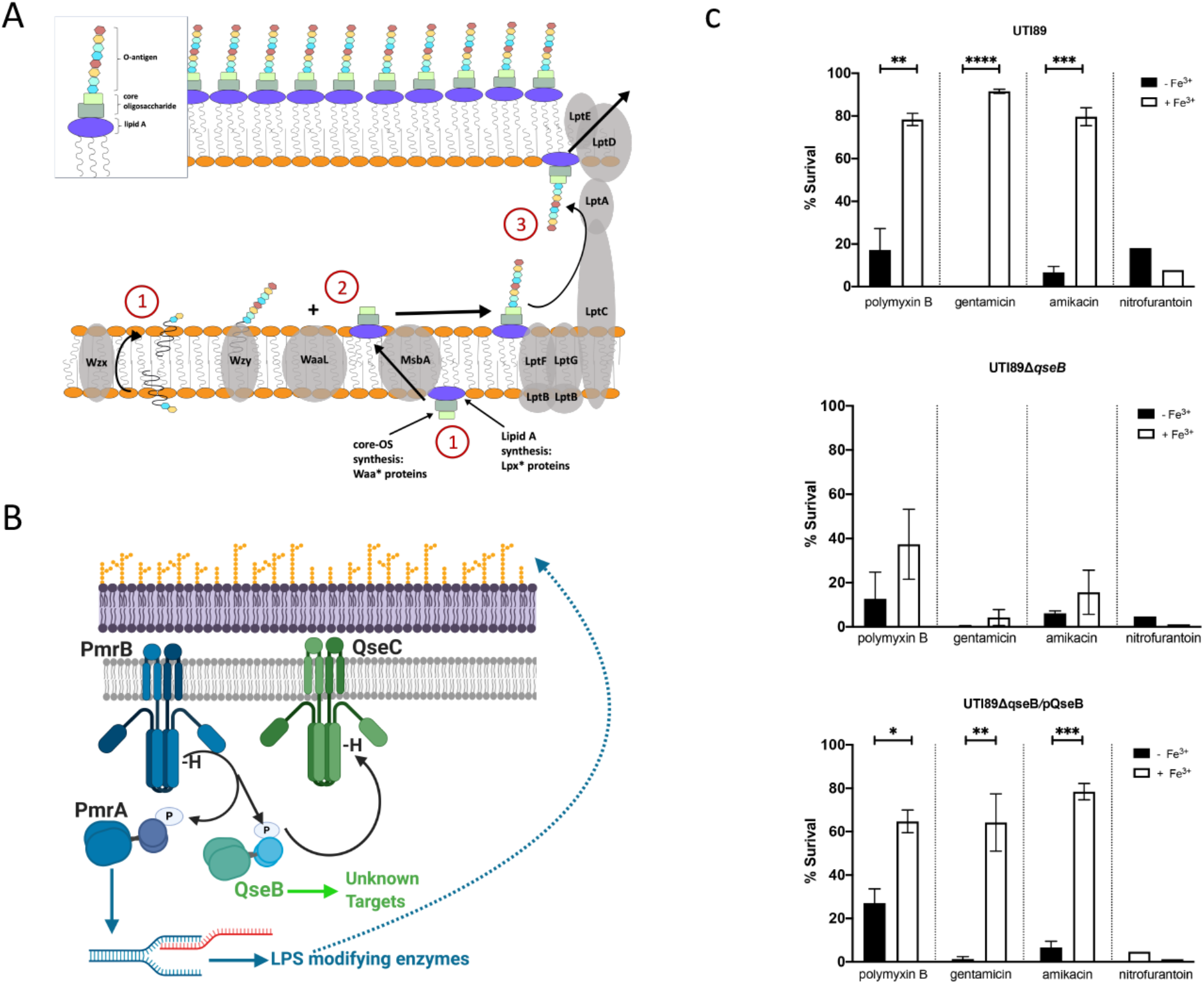
QseBC-PmrAB mediated control of polymyxin B resistance in *E. coli*. (A) Cartoon depicts - in a simplified manner - the steps in lipopolysaccharide (LPS) biosynthesis in *Escherichia coli*. (B) Cartoon depicts the mechanism of activation of the two-component systems PmrAB (blue) and QseBC (green). PmrB is a membrane bound histidine kinase that is activated by ferric iron. Upon activation it auto-phosphorylates and then transfers the phosphoryl group onto its cognate response regulator PmrA and the non-cognate QseB. PmrA acts as a transcription factor, regulating the transcription of LPS modifying genes. QseB is also transcription factor, the targets of which during antibiotic response were unknown prior to this study. (C) Graphs depict results of polymyxin B, gentamicin, and amikacin survival assays for each strain. Cells were allowed to reach mid logarithmic growth phase in the presence or absence of ferric iron and normalized. Cells were then exposed to antibiotic or to diluent alone (sterile water), for one hour. At this time cells were serially diluted and plated to determine colony forming units per milliliter (ml). To determine percent survival, cells exposed to antibiotic were compared to isogenic untreated controls (mean ± SEM, n = 3 biological repeats). To determine statistical significance, an unpaired *t*-test was performed between the untreated strain and the same strain treated with ferric iron. **, p < 0.01; ***, p < 0.001 ****, p <0.0001.

A significant body of work identified PmrA as the transcriptional activator of genes required for modifying the nascent LPS (9, 15-18). However, little is known about what controls associated metabolic shifts. Here we demonstrate that the QseB transcription factor fills in the role of metabolic controller, during UPEC’s response to polymyxin B.

We show here – for the first time through analysis of LPS modification in strains lacking different permutations of *pmr* and *qse* genes demonstrates – that QseB-PmrB signaling can result in some modification of the LPS, but not to the extent of PmrA. Transcriptional analysis of mutants deleted for either *qseB*, or *pmrA* and *qseB*, reveal that both transcription factors influence the expression of LPS modifying genes. However, deletion of *qseB* most profoundly affects metabolism genes centered on glutamate and 2-oxoglutarate-glutamate homeostasis. Accordingly, deletion of *qseB* alters glutamate levels in the cell, coincident with increased antibiotic susceptibility in the *qseB* deletion mutant. Deletion of representative QseB-regulated metabolism genes influence corresponding metabolite levels during antibiotic challenge and display a susceptibility profile that phenocopies the *qseB* deletion mutant. Exogenous addition of oxoglutarate, but not glutamate rescues the *qseB* deletion phenotype, suggesting that the cell relies on *de novo* replenishment of glutamate during the antibiotic response. Finally, analysis of clinical isolates with naturally-occurring polymyxin B-resistant subpopulations and analyses of independent in vitro evolution experiments on homogeneously polymyxin-susceptible strains reveals stable mutations in QseB-regulated targets associated with glutamate metabolism.

## MATERIALS AND METHODS

### Biological Resources: Bacterial Strains, Plasmids, and Growth Conditions

Bacterial strains, plasmids and primers used in this study are listed in Table S1. Overnight growth was always performed in liquid culture in Lysogeny Broth (Fisher) at 37°C with shaking, with appropriate antibiotics, as noted in the results. Details pertaining to growth conditions for each assay used in the study can be found in the relevant sections below.

### RNA Isolation

RNA from cell pellets was extracted using the RNeasy kit (Sigma Aldrich) and quantified using Agilent Technology (Agilent). A total of 3 micrograms (µg) of RNA was DNAse treated using the DNAfree kit (Ambion) as we previously described (19-21). A total of 1µg of DNAse-treated RNA was subjected to reverse-transcription using SuperScript III Reverse Transcriptase (Invitrogen/ThermoFisher Scientific) and following the manufacturer’s protocol.

### RNA Sequencing and Analysis

Strains were grown in N-minimal media at 37 °C with shaking, and samples were obtained as described for the transcriptional surge experiments. RNA was extracted and DNAse-treated as described in the RNA isolation section. DNA-free RNA quality and abundance were analyzed using a Qubit fluorimeter and Agilent Bioanalyzer. RNA with an integrity score higher than 7 was utilized for library preparation at the Vanderbilt Technologies for Advance Genomics (VANTAGE) core. Specifically, mRNA enrichment was achieved using the Ribo-Zero Kit (Illumina) and libraries were constructed using the Illumina Tru-seq stranded mRNA sample prep kit. Sequencing was performed at Single Read 50 HT bp on an Illumina Hi Seq2500. Samples from three biological repeats were treated and analyzed. Gene expression changes in a given strain as a function of time (15 minutes post stimulation versus unstimulated; 60 minutes post stimulation versus unstimulated) were determined using Rockhopper software hosted on PATRIC database.

### chIP-on-chip

To determine promoters bound by QseB, the strain UTI89Δ*qseB* was complemented with a construct expressing a Myc-His-tagged QseB under an arabinose-inducible promoter (22-24). As a control for non-specific pull-downs, an isogenic strain harboring the pBAD-MycHis A empty vector was used. Cultures were grown in Lysogeny Broth in the presence of 0.02µM arabinose to ensure constant expression of QseB, at concentrations similar to those we previously published as sufficient for QseBC complementation (22). Formaldehyde was added to 1% final concentration, following the methodology as described by Mooney et al., (25). Upon addition of formaldehyde, shaking was continued for 5 min before quenching with glycine. Cells were harvested, washed with PBS, and stored at -80 °C prior to analyses. Cells were sonicated and digested with nuclease and RNase A before immunoprecipitation. Immunoprecipitation was performed using an anti-Myc antibody (ThermoFisher, (23)) on six separate reactions, three for the experimental and three for the control strain. The ChIP DNA sample was amplified by ligation-mediated PCR to yield >4 μg of DNA, pooled with two other independent samples, labeled with Cy3 and Cy5 fluorescent dyes (one for the ChIP sample and one for a control input sample) and hybridized to UTI89-specific Affymetrix chips (21).

### Polymyxin B Survival Assays

To assess susceptibility of strains to polymyxin B, strains were grown in N-minimal media in the absence (unstimulated) and presence (stimulated) of ferric iron (at a final concentration of 100µM) as described for the transcriptional surge experiments and in figure 2. When bacteria reached an OD_600_ of 0.5, they were normalized to an OD_600_ of 0.5 in 5ml of 1X phosphate buffered saline (PBS) and split into two groups: A) Nothing added – “Total CFU’s control”; B) PMB added at a final concentration of (2.5 µg/mL) – “– PMB treated”. The “stimulated (+Ferric iron) samples also received ferric iron at a final concentration 100µM. Samples were incubated for 60 minutes at 37 °C after which samples were serially diluted and plated on nutrient agar plates (Lysogeny Broth agar) to determine colony forming units per milliliter (CFU/mL). Percent survival as a function of ferric iron pre-stimulation was determined by using the formula

**Figure 2:**
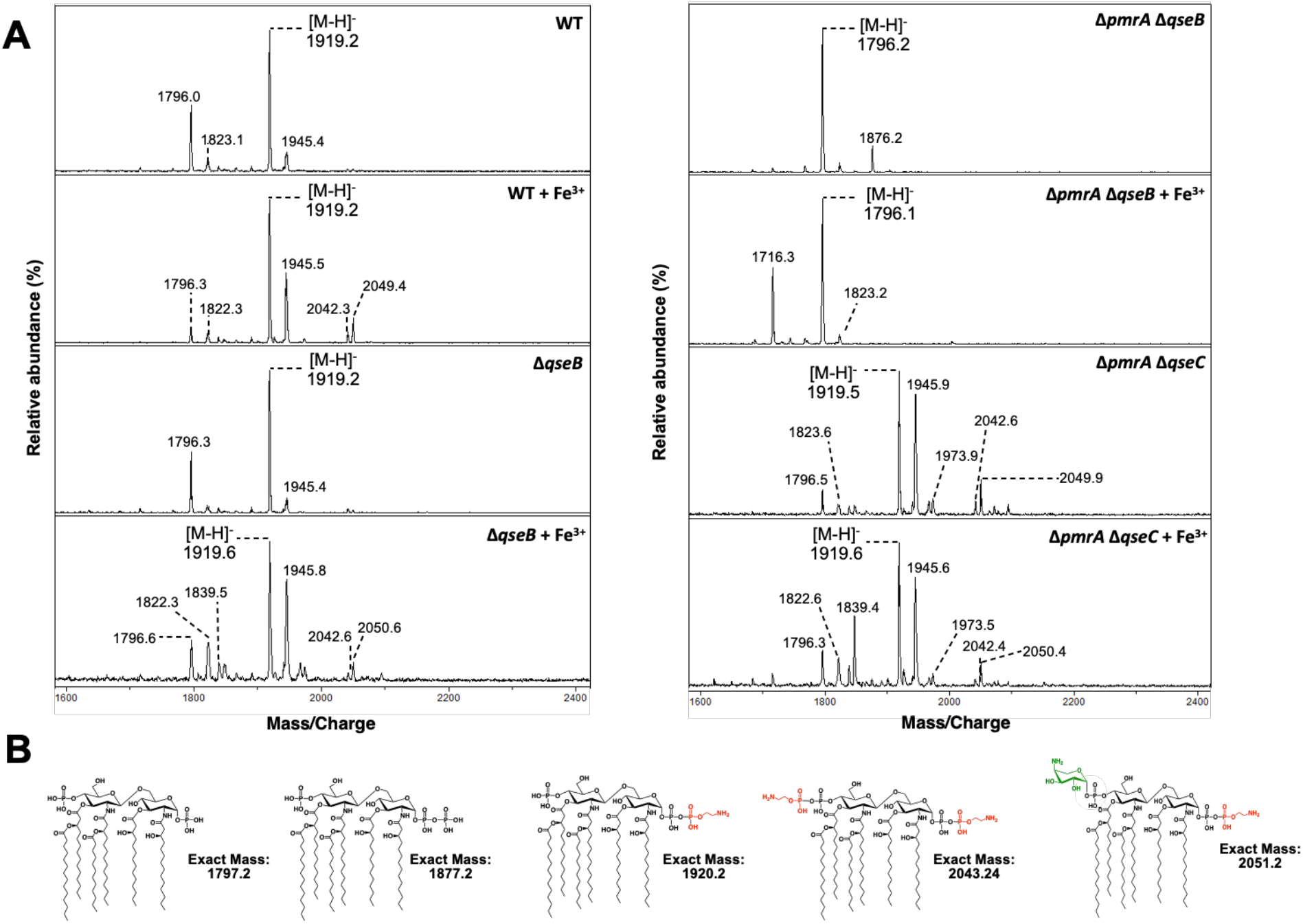
Lipid modifications in *qse/pmr* mutants. **A)** Lipid A was isolated from the indicated strains grown in N-minimal media supplemented with 10 µg/mL of niacin and iron where indicated. Lipid A was analyzed using MALDI-TOF mass spectrometry in the negative-ion mode. In UTI89 (left panel) there was unmodified (*m/z* 1796.0) and pEtN modified (*m/z* 1919.2) lipid A. For UTI89 + Fe^3+^, additional peaks were observed at 2042.3 (2 pEtNs) and 2049.4 (pEtN, L-Ara4N) representing doubly modified species. Compared to wild-type UT189, loss of *qseB* (left panel) had no effect on lipid A structure, regardless of the addition of Fe^3+^. Modification with pEtN and L-Ara4N was lost in Δ*pmrA* Δ*qseB* (+/-Fe^3+^) (right panel) with unmodified lipid A (*m/z* 1796.2) the major species. However, single and double modifications were easily detected in Δ*pmrA* Δ*qseC* (+/-Fe^3+^). Description of minor peaks: Peaks at *m/z* of ∼1822 and 1945 correspond to species detected at *m/z* of ∼1796 and 1919 containing one acyl chain extended by two carbons, respectively. The minor peak at *m/z* 1839.5 in Δ*qseB* (+Fe^3+^) contains a single pEtN, but lacks the 1-phosphate group that is easily hydrolyzed during mass spectrometry. Similarly, the peak at *m/z* 1716.3 in *pmrA, qseB* (+Fe^3+^) is the loss of 1-phosphate from unmodified lipid A. The species at *m/z* 1876.2 represents a lipid A containing a 1-diphosphate moiety giving a *tris-*phosphorylated lipid A, a species detected in the absence of activated PmrA. Data is representative of three biological experiments. **B)** Proposed chemical structures and exact masses of relevant lipid A species.

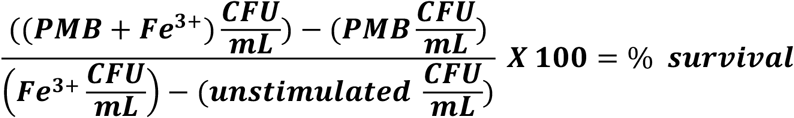

For polymyxin B (PMB), survival assays performed concurrently with metabolite measurements (see relevant section below), samples were taken across time (at induction (t=0), 15, 60, and 180 minutes post ferric iron additions). For PMB survival assays performed concurrently with oxoglutarate rescue assays (see relevant section below), oxoglutarate was added at the same time as polymyxin B after standardization of the samples to an OD_600_ = 0.5.

### Metabolite Measurements

Pellets of approximately 10^8^ cells were collected from PMB survival assays at each time-point. Glutamate, aspartate and coenzyme A levels were quantified using a colorimetric assay utilized from Glutamate-, Aspartate- and Coenzyme A Assay Kits (all kits obtained from Sigma Aldrich) utilizing the entire, undiluted sample (10^8^ cells). Assays were performed according to manufacturer’s instructions in at least 2 biological replicates per strain, per timepoint.

### Polymyxin B Minimum Inhibitory Concentration (MIC) Determination

To determine the minimum inhibitory concentration of PMB in strains used in this study the broth microdilution method was used. Strains were grown at 37°C overnight with shaking, in cation-adjusted Mueller Hinton broth following clinical microbiology laboratory standard operating procedures (26). Specifically, strains were sub-cultured at a starting OD_600_ of 0.05 and allowed to reach growth at an OD_600_ = 0.4 – 0.5. Cells were then normalized to and OD_600_ = 0.5 or roughly 10^5^ cells. At this time, a 96 well polypropylene plate was prepared with a gradient of PMB concentrations (2-fold dilution from 64 µg/mL to 0.125 µg/mL) across the rows, plus a column with no PMB added as a growth control, and a media only column to serve as a negative control. Five microliters of the standardized culture were added to each well except those holding the media control. Plates were incubated statically at 37°C for 24 hours. At this time, the minimum inhibitory concertation was determined. Minimum inhibitory concentration was set as the concentration of polymyxin B in the well in which bacterial growth was diminished by greater than 95%. Each strain was tested with 3 technical replicates and 3 biological replicates.

### Transcriptional Surge Experiments

To assess induction of *qseBC*, bacteria were grown in N-minimal media (23, 27) at 37°C with shaking (220 rotations per minute). N-minimal media were inoculated with strains of interest at starting optical density at a wavelength of 600 nm (OD_600_) of 0.05. Strains were allowed to reach mid-logarithmic growth phase (OD_600_ = 0.5). At this time, 4-milliliters of culture was withdrawn for processing (see below) and the remainder of the culture was split into two. To one of the two split cultures, ferric chloride (Fisher) was added at a final concentration of 100 µM, while the other culture served as the unstimulated control. Cultures were returned to 37°C with shaking. Four-milliliter samples were withdrawn from each culture at 15- and 60-minutes post stimulation for RNA processing, antibiotic susceptibility profiling and metabolomics as described above. All samples were centrifuged at 4000 x g for 10 minutes upon collection. The supernatant was decanted and the fraction containing the cell pellet was flash frozen in dry ice – ethanol and stored at -80°C until RNA extraction.

### Mass spectrometry of lipid A species

200 mL cultures of each strain were grown at 37°C in N-minimal media supplemented with 10 µg/mL Niacin. At an OD_600_ of ∼0.5, 100 µM of iron was added to the indicated strains and then all strains continued growing at 37°C for an additional hour. Cultures were harvested and lipid A was isolated from cells as previously described (28, 29). Mass spectra of purified lipid A were acquired in the negative-ion linear mode using a matrix-assisted laser desorption-ionization time-of-flight (MALDI-TOF) mass spectrometer (Bruker Auto-flex speed). The matrix used was a saturated solution of 6-aza-2-thiothymine in 50% acetonitrile and 10% tribasic ammonium citrate (9:1, v/v). Sample and plate preparation were done as previously described (28, 29).

### Statistical Analyses

For antibiotic survival assays, the percent survival of strains in specific conditions were calculated (mean ± SEM, N =3) and were compared to a control strain using an unpaired T-test to performed using Prism software. For polymyxin B survival assays in which several strains were compared to UTI89 with ferric iron added, a one-way ANOVA was performed with multiple comparisons. For antibiotic survival assays in which several strains were compared one another a one-way ANOVA with multiple comparisons was used. For transcriptional surge experiments across time, no statistical test was used, but the mean ± SEM was displayed. For RNA sequencing data, q-value was calculated by the Rockhopper software when calculating for differential expression between two conditions. For metabolite measurements, no statistical test was used, a representative of three biological replicates was displayed.

### Data and Code Availability

RNA sequencing data submission can be found on ArrayExpress at E-MTAB-9277. ChIP-on-chip data can be found in supplementary file S2 and is pending submission at ArrayExpress.

## RESULTS

### QseB mediates resistance to positively charged antibiotics

In previous work we determined that the PmrB sensor histidine kinase phosphorylates and activates a non-cognate response regulator, QseB that forms a two-component system with the QseC sensor histidine kinase (**Figure 1B** and (20, 22, 24)). Activation of PmrB by one of its ligands – ferric iron – leads to phosphorylation of both the cognate PmrA and the non-cognate QseB and both phosphorylation events are necessary for *E. coli* to mount transient tolerance to polymyxin B (24). Deletion of *pmrB* abolishes the ability of *E. coli* to survive polymyxin intoxication; deletion of either *pmrA* or *qseB* leads to a two-to ten-fold reduction in survival, with the double deletion mutant Δ*pmrA*Δ*qseB* phenocopying the *pmrB* deletion (**Figure S1** and (24)). While PmrA regulates the expression of LPS-modifying enzymes in both *Salmonella* and *E. coli* (15, 16), the role of QseB in the polymyxin B response in *E. coli* has been elusive. Intriguingly, studies have reported the presence of an additional *qseBC* locus within an *mcr*-containing plasmid in an isolate that is resistant to colistin, another polycationic antimicrobial, (30) further suggesting a role for the QseBC two-component system in LPS modification.

If QseB-mediated control facilitates modification of the cell envelope to a less negatively charged state, one would predict that QseB activation would also lead to tolerance to other positively charged antibiotics. To test this hypothesis, isogenic strains lacking QseB (UTI89Δ*qseB*), or carrying QseB in the native locus (UTI89) or extra-chromosomally (UTI89Δ*qseB* /pQseB), were tested for their ability to resist gentamicin and amikacin, aminoglycoside antibiotics that are positively charged. Nitrofurantoin, which is neutral, along with polymyxin B were used as negative and positive controls respectively (**Figure 1C**). Strains were tested for their ability to survive a concentration of antibiotic at up to 5 times the established minimum inhibitory concentration (MIC) (**Figure S2A**) after growth to mid-logarithmic growth phase in nutrient-limiting media as previously used to mimic the host environment and induce activation of the PmrAB two-component system (23, 24, 31). Bacterial survival was calculated during growth in media alone (black bars) or in media supplemented with 100µM ferric iron – the activating signal for the PmrB receptor (white bars), after 60 minutes. Strains harboring QseB (wild-type strain or the D*qseB*/pQseB complemented strain) exhibited 75-95% survival in the presence of positively charged antibiotics when pre-conditioned with ferric iron (**Figure 1C**, top and bottom panels). However, the strain lacking *qseB* (Δ*qseB*, **Figure 1C**, middle panel) exhibited a marked decline in survival regardless of the presence of ferric iron that was not significantly different from the survival of the un-stimulated strain. The uncharged antibiotic nitrofurantoin led to effective bacterial killing of all genetic backgrounds (**Figure 1C**). These data indicate that the QseB transcription factor mediates resistance to positively charged antibiotics.

Given the genomic plasticity associated with the species *E. coli* (32), we next asked whether QseB signaling is similar in isolates from three of the most prevalent *E. coli* phylogenetic clades, B1, B2 and E. Using a representative panel of *E. coli* strains that lack plasmid-borne polymyxin B resistance determinants and are sensitive to polymyxin by standard clinical laboratory testing (**Figure S2A**), we saw a robust transcriptional surge, an increase in transcript abundance soon after activation, followed by a slow reset, of the *qseB* promoter in all tested strains in response to ferric iron (**Figure S2B**) that coincided with an elevated survival (77%-100%) in 2.5X the MIC of polymyxin B compared to untreated controls (**Figure S2C**). Deletion of *qseB* or *qseBC* in well-characterized enterohemorrhagic *E. coli* (EHEC) strains 86-24 and 87-14 led to a significant reduction in polymyxin B tolerance compared to the wild-type parent (**Figure S3**), which was rescued upon extra-chromosomal complementation with a wild-type copy of *qseB*. Notably, strain Sakai that harbors a truncated, non-functional copy of QseC, exhibits intrinsic polymyxin B tolerance (**Figure S3)**, consistent with a model in which absence of functional QseC leads to uncontrolled PmrB-to-QseB phosphotransfer and subsequent intrinsic resistance. Supporting this notion, deletion of *qseB* or the entire *qseBC* locus in this strain phenocopies the *qseB* deletion in the other EHEC isolates (**Figure S3**). Combined these results indicate that tolerance to polymyxin B is mediated by QseB in diverse *E. coli* clades. To further probe the mechanism by which QseB mediates the response to polymyxin B, we used the uropathogenic *E. coli* (UPEC) strain UTI89.

### QseB and PmrB support LPS modifications in the absence of PmrA

We previously reported a synergistic effect of QseB and PmrA in mediating resistance to polymyxin B (24), and our data here indicate a role for QseB in mediating resistance to other positively charged antibiotics. Together, these observations suggest that QseB is involved in mediating changes to the cell envelope charge. To determine whether QseB-mediated regulation is sufficient to support LPS modification, we analyzed changes to the lipid A moiety of strain UTI89 and isogenic *<u>qse</u>* and *pmr* mutants with and without ferric iron stimulation.

Analysis of lipid A from wild-type UTI89 produced molecular ions at 1796.0 and 1919.2 *m/z* corresponding to unmodified lipid A and the addition of a single pEtN, respectively (**Figure 2A**). With the addition of Fe^3+^ to the growth media, additional modifications were apparent, including lipid A with two pEtN residues (2042.3 *m/z*) and a species modified with both L-Ara4N and pEtN (2049.4 *m/z*) indicating increased levels of lipid A modification. Deletion of *qseB* in wild-type UTI89 had no impact on lipid A structure and similar molecular ions were detected in the presence (*m/z* 1919.6, 2042.6, 2050.6) or absence (*m/z* 1796.3, 1919.2) of iron (**Figure 2A**). The additional species at *m/z* 1839.5 in the *qseB* mutant with Fe^3+^ arises from the loss of the 1-phosphate that is easily hydrolyzed during mass spectrometry from the [pEtN]_2_-lipid A species (33-35). This result indicates that QseB loss does not impair the covalent modification of lipid A.

The pEtN modification was lost in the UTI89Δ*pmrA*Δ*qseB* double mutant; only unmodified lipid A (*m/z* 1796.2) was present. Furthermore, addition of Fe^3+^ could not restore lipid A modifications in UTI89Δ*pmrA*Δ*qseB*. Given the current literature, this was expected since PmrA is necessary for transcription of genes encoding lipid A modification machinery and solidify that PmrA is the primary controller of LPS modification genes. However, single and double modified species were easily detected in strain UTI89Δ*pmrA*Δ*qseC* (**Figure 2**), suggesting that QseB also regulates expression of *eptA (pmrC, yjdb) and arnT (pmrK)*. Furthermore, the addition of Fe^3+^ was no longer required for production of doubly modified species in the UTI89Δ*pmrA*Δ*qseC* consistent with our previous report that in this strain PmrB constitutively activates QseB and results in a strain that shows intermediate levels of polymyxin resistance according to CLSI standards (24). Together, these results indicate that – although PmrA is undoubltedly the primary regulator of lipid A modifications – QseB can support some lipid A modifications, suggestive of a transcriptional regulatory overlap with PmrA. Moreover, given that loss of QseB did not alter the type of lipid A modifications, these data suggest that QseB must play a distinct role in the modulation of antibiotic tolerance.

### RNAseq profiling reveals regulatory redundancy between PmrA and QseB

The lipid A profiling suggests that QseB regulates *eptA and arnT* expression, since these are the enzymes responsible for the observed modifications in UTI89Δ*pmrA*Δ*qseC*. To decipher whether there are regulatory overlaps in transcription patterns between QseB and PmrA, steady-state transcript abundance across the activation surge were tracked over time via RNA sequencing (RNAseq). For these RNAseq experiments, the wild-type strain UTI89 and isogenic UTI89Δ*qseB*, UTI89Δ*pmrA*Δq*seB* or UTI89Δ*pmrA*Δq*seC* were grown under PmrB-activating conditions (100µM Ferric iron, Fe^3+^) and samples were obtained for RNA sequencing immediately prior to (T=0), as well as 15 (T=15) and 60 (T=60) minutes post addition of ferric iron to the growth medium (**Figure 3A**). Output RNA sequencing data from three biological repeats per strain per timepoint were analyzed using Rockhopper software (30, 31). Differential gene expression matrices within each strain were calculated to compare T = 0 to T =60 and T =15 minutes (**Figure 3, Supplementary File 1**). An additional comparison of T=15 and T=60 was also made (**Supplementary File 1**). Pairwise differences *across* strains were analyzed for each timepoint (**Supplementary File 2**). Transcripts with a q value lower than 0.05 were considered significant (**Supplementary Files 1-2**). These analyses demonstrated that in the wild-type strain LPS modification gene expression surged over time, following stimulation with ferric iron (**Figure 3B**). However, the same clusters had no significant surge in the mutants lacking QseB (**Figure 3B, Supplementary File 1**). Comparison of UTI89Δ*pmrA*Δ*qseC* to wild-type UPEC at t=0, t=15 and t=60 revealed that indeed in the UTI89Δ*pmrA*Δ*qseC* strain, both *arnT* and *eptA* (*yjdB*) are induced, along with other members of the *arn* operon (**Supplementary File 2**). Together, these data indicate that QseB and PmrA share regulatory redundancy in mediating transcription of LPS modification genes.

**Figure 3.**
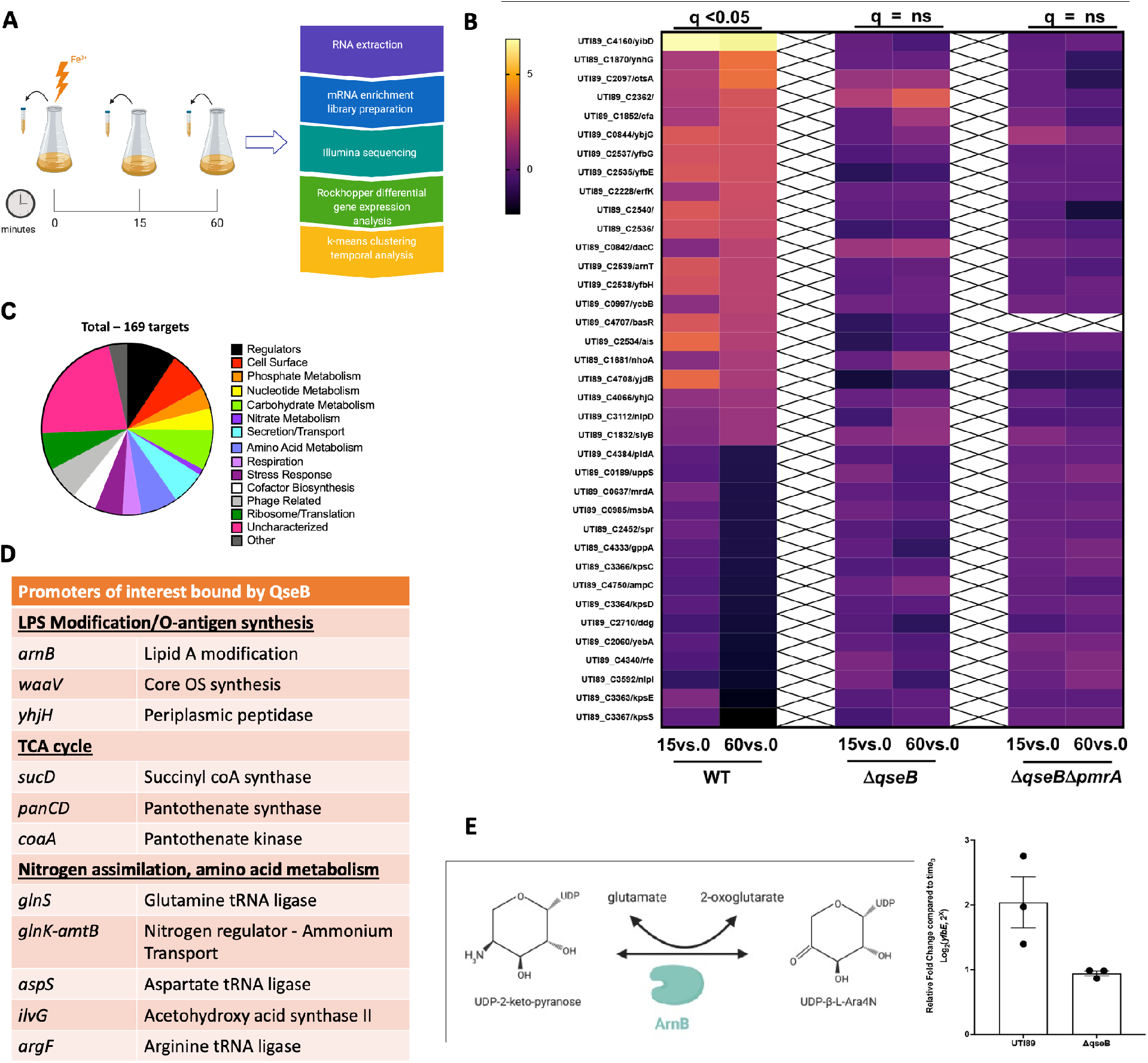
QseB and PmrA have regulatory overlaps. **A)** Schematic shows the pipeline for sample collection and data processing in the presented RNAseq study. **B**) Heatmaps indicate log_2_ relative fold change of wild-type (WT) UTI89 and the isogenic Δ*qseB*, and Δ*qseB*Δ*pmrA* strains, for genes involved in metabolism after stimulation with ferric iron at 15- and 60-minutes post stimulation. These genes were significantly (q<0.05) changed at 60 minutes compared to pre-stimulation (T=0) in wild-type UTI89, but showed no statistically significant change in the absence of *qseB* alone or in the absence of both *qseB* and *pmrA*. **C-D)** Direct targets of QseB identified in chIP-on-chip analyses. **C)** Pie chart indicates the distribution of 169 unique DNA promoter sequences bound by QseB in pull-down experiments using tagged QseB and cross-linking, followed by immunoprecipitation, reversal of the crosslinks and hybridization of eluted DNA onto Affymetrix UTI89-specific chips. The data are from three independent biological experiments and exclude non-specific targets isolated through immunoprecipitation with vector control. **(D)** Subset of the direct targets of QseB. **E)** Cartoon depicts the conversion of UDP 2-keto pyranose to UDP-β-L-Ara4N by ArnB, in a reaction that consumes a glutamate molecule and produces an oxoglutarate molecule in the process. The graph depicts the expression of *yfbE* (*arnB*) at 60 minutes post stimulation with ferric iron in UTI89 and UTI89Δ*qseB*. Briefly, cells were grown to mid-log growth phase. Cells were collected before stimulation and at 60 minutes post stimulation with ferric iron for RNA extraction and reverse transcription. Resulting cDNA was subjected to qPCR with a probe complementary to the *yfbE* region (See Table S1 for corresponding primers and probe). Graph depicts log_2_-fold change of *yfbE* transcripts at each time point relative to the sample taken before stimulation (mean ± SEM, n = 3 biological repeats, depicted as dots in the graph).

To further validate RNAseq experiments and to identify promoters bound by QseB, we performed a chiP-on-chip experiment, using the UTI89Δ*qseB* strain that harbors a construct expressing Myc-His-tagged QseB under an arabinose-inducible promoter (22-24). An isogenic strain harboring the pBAD-MycHis A vector was used as a negative control. Pull-downs using an anti-Myc antibody were performed on six separate reactions, three for the experimental and three for the control strain. Analyses of the pull-down DNA revealed a total of 169 unique promoters bound by QseB and absent in the negative control (**Figure 3C-D** and **Supplementary File 1**). Among the promoters identified was the promoter of *qseBC* **(Supplementary File 1)** – consistent with QseB’s ability to regulate its own transcription (21, 22, 36), as well as *yibD*, which we have previously validated as a QseB binding target (23, 24). Another promoter identified was indeed the *arnBCADTEF* promoter region, with portion of the *arnB* gene, which is the first gene in the operon, also pulled down the analyses (**Figure 3D, Supplementary File 1)**. Accordingly, qPCR analysis determined *arnB* transcript levels were 2.16 times lower in the *qseB* deletion mutant compared to the wild-type control **(Figure 3E**), validating that QseB transcriptionally regulates the *arn* operon.

### QseB controls central metabolism genes

Our RNAseq and ChIP-on-chip profiling revealed that in addition to LPS-modification, QseB controls several genes that code for central metabolism enzymes (**Figure 4A, Figure 3D, and Supplementary File 1**). Among the most highly upregulated genes in the RNAseq profiling were genes involved in arginine and isoleucine biosynthesis, as well as genes encoding TCA cycle enzymes (**Figure 4A and Supplementary File 1**). However, the same clusters had no significant surge in the mutants lacking QseB (**Figure 4A** and **Supplementary File 1**). The chIP-on-chip analyses revealed *ilvG* and *argF* as QseB direct targets (**Figure 3D, Supplementary File 1**) as, well as another set of metabolic targets including *glnK, glnS* and *aspS* that are involved in glutamine-glutamate and aspartate-glutamate interconversions respectively (**Figure 3D and Supplementary File 1**). In media with low ammonia, glutamate can be synthesized via the combined action of glutamine- and glutamate synthases, encoded by *glnA* and *gltBD* respectively (12). This occurs via the condensation of glutamate with ammonia by GlnA, followed by reductive transamination of the produced glutamine with oxoglutarate by GltBD. Under high nitrogen/ammonia conditions, glutamate is synthesized by glutamate dehydrogenase encoded by *gdhA* (12). In the growth conditions used in our studies, ammonia was limiting, but the overall nitrogen concentration was 7.5mM. The *glnA* and *gdhA* loci were not part of the upregulated genes, but *ybaS* – which also converts glutamine to glutamate (37) – and *gltA, gltB* and *gltD* were among the most highly upregulated genes, the surge of which depended on QseB (**Figure 4A**). Consistent with previous studies demonstrating a high ATP and NADPH requirement for nitrogen assimilation/glutamate production (13, 38), genes involved in aspartate, beta alanine oxaloacetate conversions, as well as a possible glutamate-fumarate shunt were identified as key QseB regulated targets (**Figure 4A-B**). The genes involved in glutamate metabolism are particularly intriguing, given that the *arnB* gene product catalyzes transamination of undecaprenyl-4-keto-pyranose to undecaprenyl 4-amino-4-deoxy-L-arabinose (**Figure 3E**), which consumes glutamate, releasing oxoglutarate in the process. Previous work by the Raetz group indicated that this reaction is not energetically favored (11). We thus asked whether QseB regulates glutamate-oxoglutarate homeostasis during *E. coli’s* response to positively charged antibiotics.

**Figure 4.**
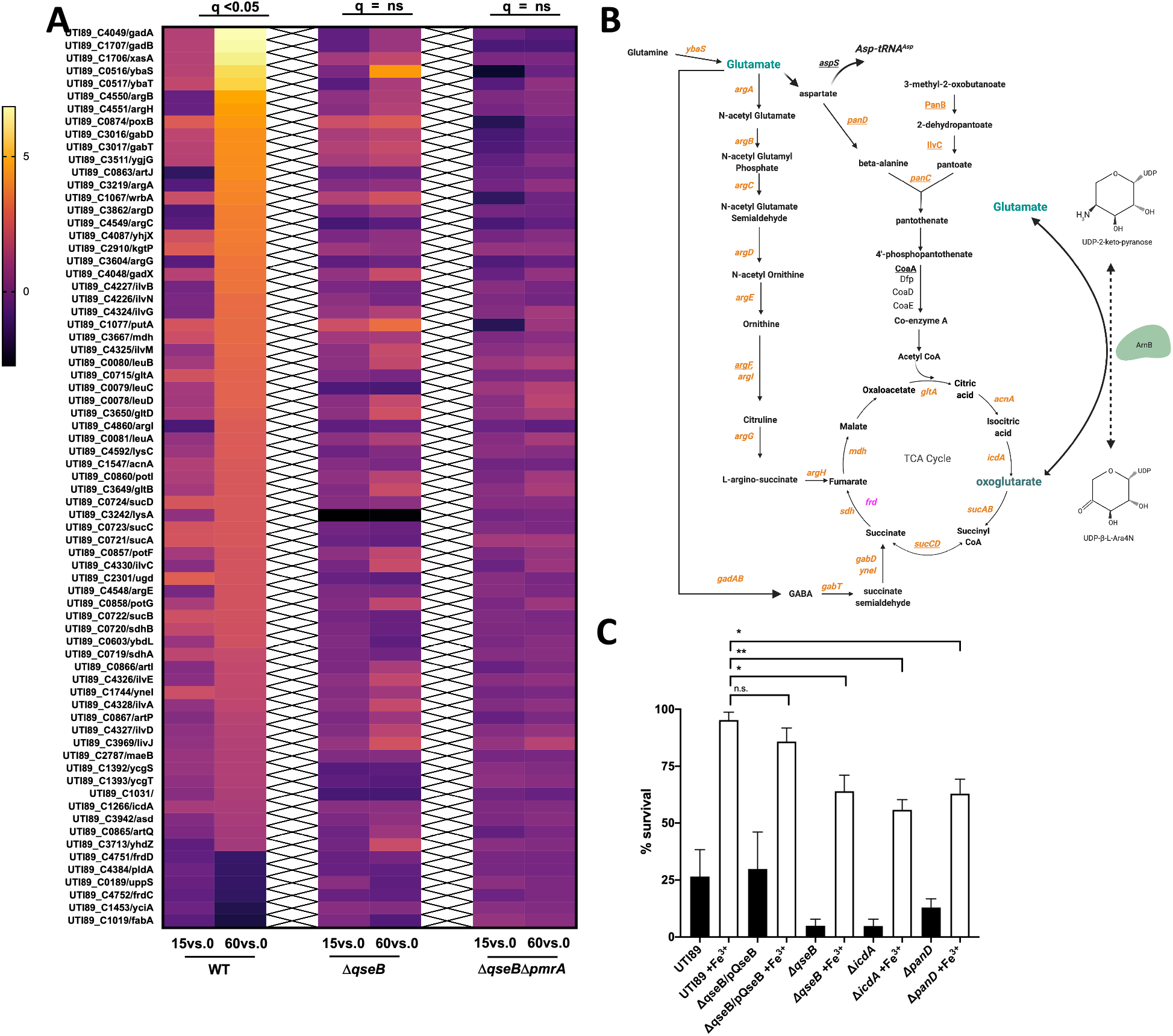
RNAseq profiling across time reveals a metabolic circuit under QseB control. **A)** Heatmaps indicate log_2_ relative fold change of wild-type (WT) UTI89 and the isogenic Δ*qseB*, and Δ*qseB*Δ*pmrA* strains, for genes involved in metabolism after stimulation with ferric iron at 15- and 60-minutes post stimulation. These genes were significantly (q<0.05) changed at 60 minutes compared to pre-stimulation (T=0) in wild-type UTI89, but showed no statistically significant change in the absence of *qseB* alone or in the absence of both *qseB* and *pmrA*. **B)** Schematic outlines the metabolic pathways regulated by QseB. Genes in orange are upregulated at 60 minutes post-stimulation with ferric iron as indicated by RNA sequencing data. Genes in pink are downregulated at 60 minutes post-stimulation with ferric iron. Genes underlined are direct targets of QseB as indicated by ChIP-on-chip data. **C)** Graph depicts polymyxin B survival assays for wild-type *E. coli* and isogenic mutants deleted for *qseB, icdA, or panD*. Cells were allowed to reach mid logarithmic growth phase in the presence or absence of ferric iron and and normalized. Cells were then either exposed to polymyxin at 2.5 μg/mL or without addition for one hour. To determine percent survival, cells exposed to polymyxin were compared to those that were not (mean ± SEM, n = 3). To determine statistical significance, a one-way ANOVA with multiple comparisons was performed between strains treated with ferric iron and UTI89 wild-type treated with ferric iron. *, p < 0.05. N.S. denotes a comparison that did not result in statistical significance.

### Glutamate – oxoglutarate homeostasis, regulated by QseB, is necessary for mounting antibiotic resistance

To determine how the QseB-regulated metabolism genes would influence antibiotic resistance in a strain that contains both ArnB and QseB, we turned to a combination of metabolomics and mutagenesis. First, we created deletions in *panD* and *icdA*. PanD codes for an aspartate decarboxylase that converts aspartate into beta-alanine, which then feeds into the pantothenate pathway eventually resulting in coenzyme A production (**Figure 4B**). The *pan* gene operon is under the direct control of QseB, as the operon’s promoter was bound by QseB (**Figure 3D and Supplementary File 1**). We reasoned that if the identified QseB regulon is active during LPS modification, then we would detect changes in coenzyme A production and that deletion of *panD*, which is centrally placed in the identified pathway (**Figure 4B**) should impair antibiotic resistance. In parallel, we created a control *icdA* deletion mutant, disrupting the conversion of isocitrate to oxoglutarate (**Figure 4B**), thereby limiting oxoglutarate production, which we reasoned would be needed for GltAB activity. Obtained mutants were tested in polymyxin B survival assays alongside the wild-type parental strain and the isogenic *qseB* deletion mutant, as well as the *qseB* deletion mutant complemented with *qseB*. Strains were tested for their ability to survive a concentration of PMB at five times the established MIC. While the wild-type and the Δ*qseB*/pQseB complemented strains exhibited 85-95% survival in 5X the PMB MIC when pre-conditioned with ferric iron, the *qseB, panD* and *icdA* deletion mutants reproducibly exhibited a 50% reduction in survival (**Figure 4C**). Metabolite measurements of aspartate and coenzyme A, which are the first and last metabolites in the identified PanD pathway (**Figure 4B**), revealed altered aspartate and coenzyme A abundance in cells devoid of QseB compared to wild-type samples (**Figure S4**), indicating, that QseB indeed influences production of these intermediates.

To determine how deletion of *qseB* influences glutamate levels during an antibiotic stress response, we quantified glutamate in wild-type UPEC, the Δ*qseB* strain and the Δ*qseB/*pQseB complemented control. For metabolomics measurements, samples were taken from wild-type UTI89, UTI89Δ*qseB* and the complemented strain UTI89Δ*qseB/*pQseB grown in the presence of ferric iron (PmrB activating signal) (**Figure 5**, red lines), polymyxin B (PMB) alone (**Figure 5**, blue lines), or in the presence of ferric iron and PMB (**Figure 5**, pink lines). Control cultures in which no additives were included (**Figure 5**, black lines) were also included. In the wild-type background, addition of polymyxin B, or polymyxin B/ferric iron, resulted in a rapid decrease in glutamate levels compared to cells exposed to ferric iron alone or no additives (**Figure 5A**). However, in the Δ*qseB* deletion strain there was no change in glutamate levels in the different growth conditions (**Figure 5B**). Complementation of Δ*qseB* with pQseB, which restores extrachromosomal expression of QseB from a high-copy plasm, results in a drop in glutamate levels shortly after addition of ferric iron, polymyxin B, or both (**Figure 5C**). These data indicate that glutamate levels change during the antibiotic response, in a manner that depends upon QseB

**Figure 5.**
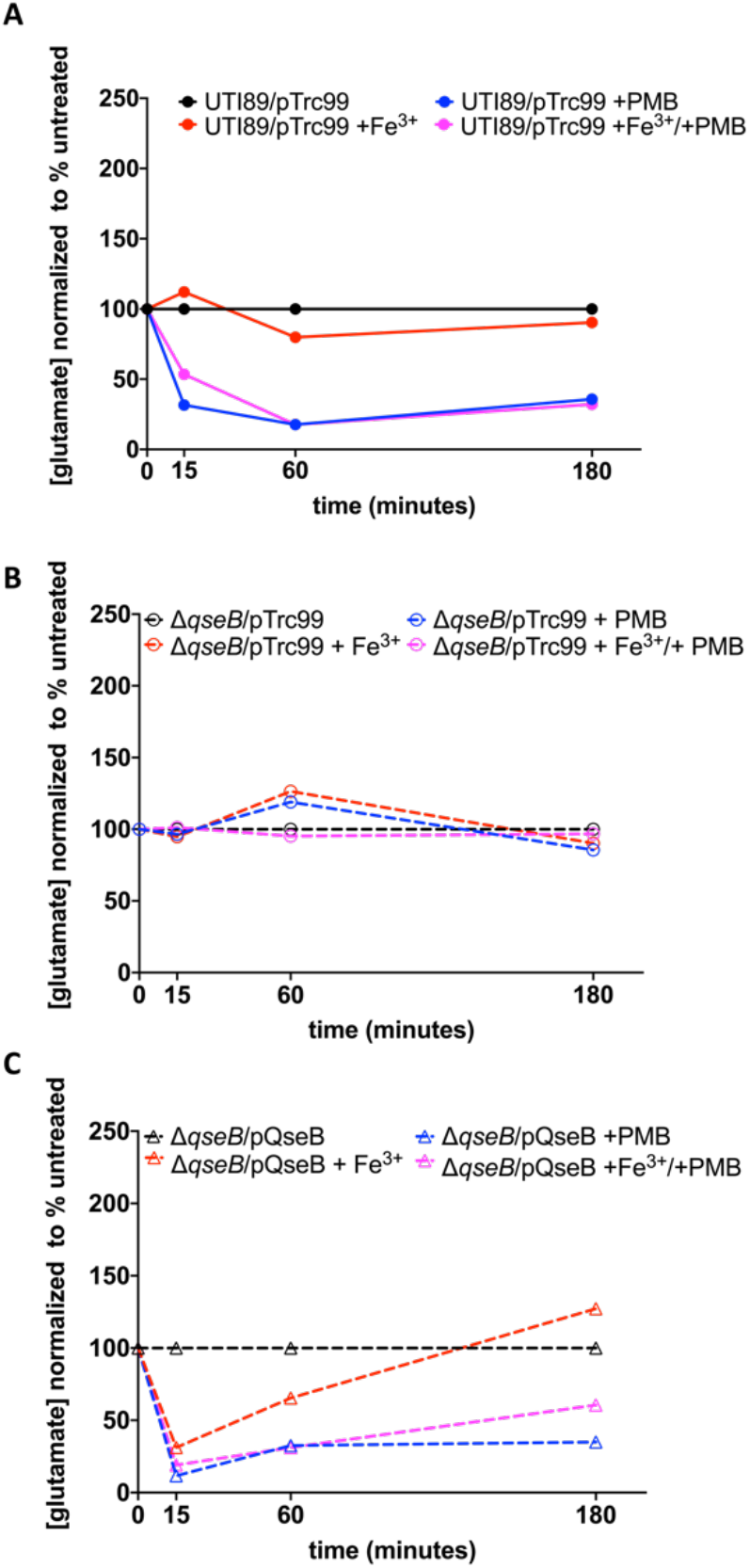
Glutamate metabolism is under the control of QseB. **A-C)** Graphs depict glutamate abundance over time in wild-type *E. coli* and isogenic mutants under different stimulation conditions. Measurements are normalized to a sample in which no additives or conditions were changed (black lines). Pink lines show measurements from samples in which ferric iron and polymyxin were added. Blue lines show measurements from samples in which only polymyxin was added. Red lines show measurements in which only ferric iron was added. Glutamate was measured across time in wild-type UTI89 (**A**), UTI89Δ*qseB*/pTrc99 (**B**) and UTI89Δ*qseB/pQseB* (**C**). A representative of three biological replicates is depicted.

If our transcriptional, metabolic and antibiotic resistance results point towards a central glutamate production circuit that is controlled by QseB and requires oxoglutarate, we next asked whether the susceptibility of the *qseB* deletion mutant to positively charged antibiotics could be rescued via the addition of exogenous oxoglutarate or glutamate. Addition of oxoglutarate to polymyxin B-treated samples of Δ*qseB* restored survival in 5X the antibiotic MIC (**Figure 6A**). Addition of glutamate did not have the same effect (**Figure 6B**). These data suggest that sufficient uptake of glutamate may not be occurring in the absence of QseB and that oxoglutarate-glutamate homeostasis controlled by QseB is necessary for mounting resistance to positively charged antibiotics.

**Figure 6.**
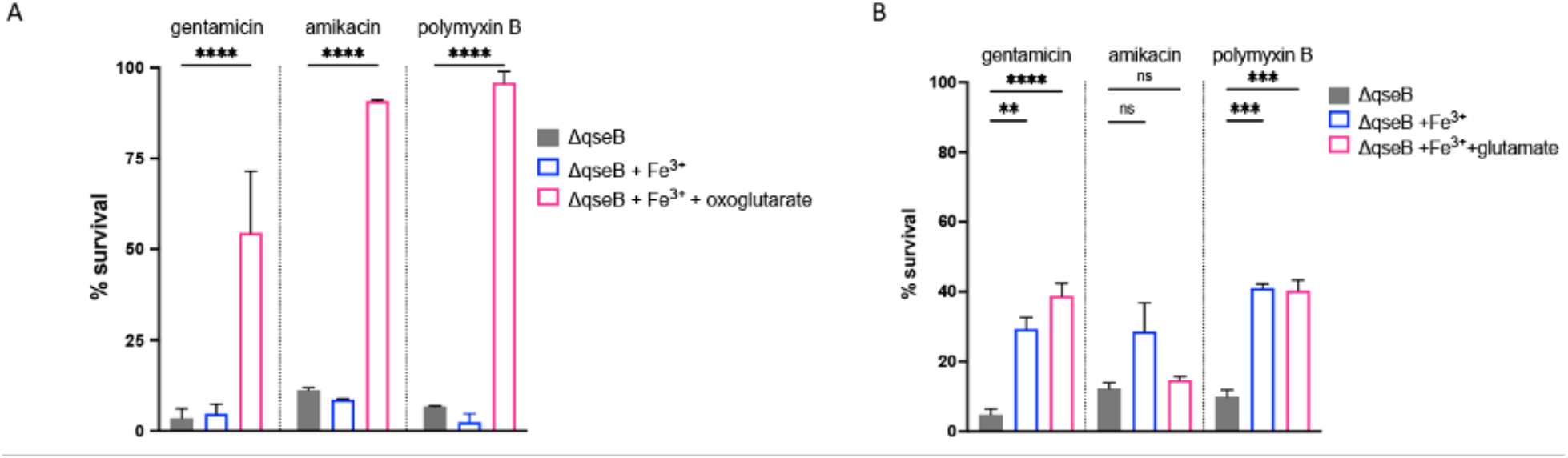
Addition of exogenous oxoglutarate rescues *qseB* deletion mutant. **A-B)** Graphs depict results of polymyxin B, gentamicin, and amikacin survival assays for the D*qseB* deletion mutant in the presence or absence of exogenous oxoglutarate (A) or glutamate (B). Cells were allowed to reach mid logarithmic growth phase in the presence or absence of ferric iron and normalized. Cells were then exposed to antibiotic or to diluent alone (sterile water), for one hour. An additional subset of cells received both ferric iron and oxoglutarate (**A**) or glutamate (**B**). At this time cells were serially diluted and plated to determine colony forming units per milliliter. To determine percent survival, antibiotic-treated cells in which metabolite was added were compared to the antibiotic-treated controls that were not supplemented with oxoglutarate or glutamate (mean ± SEM, n = 3 biological repeats). To determine statistical significance, a one-way ANOVA was performed with multiple comparisons between the untreated- and treated samples. **, p < 0.01; ***, p < 0.001, N.S, no statistical significance detected by test used.

## Discussion

Bacteria can mount resistance to antibiotics through acquisition of mobile genetic elements, including plasmids that code for antibiotic resistance cassettes. However, in many pathogens, resistance to antimicrobial agents is encoded chromosomally. In *E. coli* and *Salmonella spp*., resistance to positively charged antibiotics is intrinsically encoded in LPS modification genes. Yet, this intrinsic mechanism comes at a metabolic cost associated with diverting central metabolites to synthesize modified LPS. Our work builds upon this model and begins to unravel the complex metabolic consequences of antibiotic resistance.

While numerous studies have extensively described the various enzymes and pathways that contribute to antibiotic resistance, few have evaluated the metabolic impact of these chemical reactions on the cell. Moreover, the studies that do evaluate the metabolic impact of antibiotic resistance have primarily focused on global metrics such as population growth rate tradeoffs. More recently, several groups have begun to evaluate the influence of central metabolism on antibiotic susceptibility and are converging on a model whereby central metabolic activity – including respiratory rate and TCA cycle flux – plays a determining factor in antibiotic susceptibility (39-43). Bactericidal antibiotics can exert toxic effects on the cell by elevating metabolic rate and promoting ROS production, and these effects can be mitigated by reducing metabolic activity. In addition to affecting the overall population growth and metabolic rates, antibiotic resistance mechanisms often consume central metabolites and accordingly exert a significant effect on cellular metabolism. In this work, we demonstrate that the generation of transiently antibiotic resistant bacteria leads to a wholesale rewiring of central metabolism that may allow the cell to compensate for the consumption of metabolites during the reactions that generate antibiotic resistance.

We make two significant contributions to the field: 1) We demonstrate that in *E. coli*, QseB and PmrA share common targets in the genes that modify the LPS, but QseB plays a unique role in controlling central metabolism during LPS modification. 2) We demonstrate – through the requirement for oxoglutarate – that the anaplerotic routes identified in our analyses feed back into the TCA cycle to elevate metabolic rate and balance the glutamate necessary for mounting resistance to this class of antibiotics, without jeopardizing the cell’s ability to assimilate nitrogen, a process that largely depends on glutamate (12). By upregulating these pathways, the cell may compensate for the metabolic consequences of antibiotic resistance by regenerating and rebalancing the concentration of critical reaction intermediates.

QseB directly targets and controls several genes involved in fueling the TCA cycle during the response to positively charged antibiotics. Our data suggest that increased production of oxoglutarate by the modification of the lipid A domain of LPS may increase flux through the TCA cycle that is fueled – at least in part – by QseB-regulated pathways. Intriguingly, this process also requires the consumption of glutamate, which may be in part restored by the reversible reaction of ArnB. Our metabolomic data point towards the use of glutamate through the pantothenate pathway and co-enzyme A production, which can then enter the TCA cycle either as acetyl-CoA or succinyl-CoA (**Figure 4B**). Likewise, conversion of glutamate to fumarate via the *arg* gene products would supply fumarate, while conversion of glutamate to GABA through the function of the *gab/gad* would re-introduce succinate into the TCA cycle. This step would replenish succinate, bypassing the need to convert oxoglutarate to succinate via the *sucAB-* and *sucDC-*encoded complexes (**Figure 4B**). This could divert oxoglutarate to produce glutamate via the activity of GdhA. Future work will focus on delineating the effects of QseB on GABA abundance and GdhA-mediated production of glutamate.

In studying emerging antibiotic resistance mechanisms, we tend to generally focus on understanding plasmid-encoded systems. Here, understanding a chromosomally encoded system may translate to emerging plasmid encoded systems, and pose a threat in the clinic, given that a new plasmid, *mcr*-9, encoding both a *mcr-*1 homologue and *qseBC*-like elements has been recently reported (30). This is especially concerning, because cationic antimicrobials, such as colistin are considered “antibiotics of last resort” and reserved for multi-drug resistant infections. The finding that *E. coli* and potentially other Enterobacteriaceae have the potential to mount an intrinsic resistance response to polymyxins and aminoglycosides raises the alarm for the need to better understand mechanisms that lead to heterogeneous induction of systems like QseBC in the bacterial pathogens. Lastly, this work demonstrates the need to understand how metabolic pathways can be exploited in pathogenic bacteria and may give new insights to potential therapeutic targets.

## Acknowledgements

The authors would like to acknowledge the Center for Innovative Technologies for providing metabolomics support; the laboratory of Dr. Scott J. Hultgren for supporting the chIP-on-chip experiments through the following sources of funding: P50 DK64540, R01 AI048689 and R01AI02549; Dr. Erin J. Breland for helpful discussions regarding the transcriptional profiling; and the following sources of funding: R01 AI 5R01AI107052 and 1P20DK123967-01 to MH, R01 AI129940, R01 AI138576, R01 AI150098 to MST. Melanie Hurst is supported by NRSA F31 fellowship 1F31AI143244-01A1.

## Author Contributions

MNH, CJB and MH designed, executed and interpreted experiments, prepared figures and wrote the manuscript. KRG designed and performed RNAseq experiments and edited the manuscript. AP and MST performed the LPS modification analyses. TB, AH, SAR and DW performed antibiotic resistance experiments, analyzed RNAseq profiling data (blinded) and edited the manuscript.

